# Hidden cell diversity in Placozoa: Ultrastructural Insights from *Hoilungia hongkongensis*

**DOI:** 10.1101/2020.11.13.382382

**Authors:** Daria Y. Romanova, Frederique Varoqueaux, Jean Daraspe, Mikhail A. Nikitin, Michael Eitel, Dirk Fasshauer, Leonid L. Moroz

## Abstract

From a morphological point of view, placozoans are among the most simple free-living animals. This enigmatic phylum is critical for our understanding of the evolution of animals and their cell types. Their millimeter-sized, disc-like bodies consist of only three cell layers that are shaped by roughly six major cell types. Placozoans lack muscle cells and neurons but are able to move using their ciliated lower surface and take up food in a highly coordinated manner. Intriguingly, the genome of *Trichoplax adhaerens*, the founding member of the enigmatic phylum, has disclosed a surprising level of genetic complexity. Moreover, recent molecular and functional investigations have uncovered a much larger, so-far hidden cell-type diversity. Here, we have extended the microanatomical characterization of a recently described placozoan species – *Hoilungia hongkongensis*. In *H. hongkongensis*, we recognized the established canonical three-layered placozoan body plan but also came across several morphologically distinct and potentially novel cell types, among them novel gland cells and “shiny spheres”-bearing cells at the upper epithelium. Thus, the diversity of cell types in placozoans is indeed higher than anticipated.

## Introduction

Placozoans are millimeter-sized disc-shaped marine animals without organs, muscles or neurons. In the last decades, alongside other aquatic invertebrates, placozoans have become important model organisms to understand the origins and evolution of animal cell types, including the rise of neuronal communication^1–4^.

The first placozoan, *Trichoplax adhaerens* – an “adhering hairy plate” – was discovered in a seawater aquarium in Graz, Austria, towards the end of the 19th century^5^ (Schulze, 1883) (for a brief history of the discovery of *T. adhaerens* see^6^). In a very careful histological study, Franz Eilhard Schulze observed under the light microscope that the flat animal had a highly changeable shape and lacked symmetry^7^, and that only top vs. bottom and marginal vs. interior can be distinguished. As he could not place the animal in any known animal phyla, he tentatively suggested that *T. adhaerens* could be a very basal metazoan.

Schulze found that the animal is moving with the help of cilia at its lower surface. He also described that the animal propagates by fission. The flat body of *T. adhaerens* is organized into just three thin cell layers, an upper (water-facing) squamous epithelium and a lower (surface-contacting) columnar epithelium; in-between the two ciliated epithelia, he observed a loose layer of possibly contractile spindle- or star-like cells. Among other things, he reported that the upper side contains large refractive inclusions, so-called “shiny spheres” (“Glanzkugeln”), while he saw smaller “matte-finished spheres” (“mattglänzende Kugeln”) embedded in the lower epithelium, where the animal takes up its food. Note that the animal’s upper and lower sides have been described as dorsal and ventral in earlier publications. To prevent a currently unsupported homologization with the bilaterian dorsal/ventral axis, we refrain from using these terms when referring to the top-bottom axis of placozoans.

It was only in the early 1970s that Karl Grell and others started providing the first detailed electron microscopic characterization of *T. adhaerens*^8–14^. They confirmed Schulze’s earlier results that *T. adhaerens* is composed of three thin cell layers, formed by four different cell types: The flat and T-shaped upper epithelial cells (i), which have a single cilium; embedded in the upper surface, they reported large granules, corresponding to the “shiny spheres”. The cylindrical cells of the lower epithelium (ii) were found to bear one cilium surrounded by microvilli-like structures. Between these lower epithelial cells, sporadic “gland cells” (iii) without cilia were described. Note that “gland cells” since turned as a generic term used to refer to cells that appear to contain a large number of secretory vesicles, independently of the presence/absence of a cilium. In the epithelial interspace, they observed elongated fiber cells (iv), which form a network and can contact other cell types.

Grell also reported that the animal grazes on algae. For this, it moves over the algae, releases digestive enzymes, and takes up the predigested food at the lower epithelium. Upon this careful microanatomical analysis, Grell placed *T. adhaerens* into its own phylum, called Placozoa^15^.

In 2014, the morphology of *T. adhaerens* was revisited upon preservation in a living-like state using high-pressure freezing of the live animal combined with electron microscopy^16^. Two novel cell types were described: lipophil cells (v) and crystal cells (vi). Lipophil cells are sitting in the lower epithelium and extend deep into the interior. They often possess a large vacuolar structure at their distal end. It is likely that these distal vacuole-like structures correspond to the matte-finished spheres described by Schulze^7^. These cells are fragile and probably had not been preserved during chemical fixations carried out by Grell and others. Crystal cells contain birefringent crystals that have been reported earlier^17^. Those crystals are made of aragonite^18^, which are thought to serve as gravity sensors and contribute to geotaxis^18^. In another EM study, small ovoidal cells (vii) have been observed in the marginal zone, where the two epithelia meet^19^.

A recent single-cell transcriptome study has revealed a hidden cell type diversity in *Trichoplax*^2,3^. This study did not identify the distinct expression profile of a putative muscle or neuronal cell, corroborating the morphological notion that placozoans lack such specialized cell types. However, profiles of specialized cells involved in digestion and metabolism and several cells expressing hormones and peptides were readily identified^3^. The much larger cell diversity was corroborated by studies that investigated the function of the surprisingly diverse repertoire of small peptide hormones in *T. adhaerens*. Different, non-overlapping populations of peptide-releasing cells exist in fixed locations in all three layers of the animal^20,21^. In addition, other studies of Smith and coworkers revealed the presence of additional cell types in *T. adhaerens*, including mucus-releasing cells, and distinguishing two types of ciliated gland cells in the lower epithelium^22^.

The first specimens of *T. adhaerens* (also known as mitochondrial 16S haplotype H1) originated from the Mediterranean and the Red Sea; this species was used in the majority of placozoan studies. However, broad biodiversity surveys performed during the last decade identified dozens of 16S haplotypes^23–30^. The ecological niche occupied by placozoans covers the sublittoral zone of tropical, subtropical, and some temperate waters from 0.5 to 20 meters deep and potentially contains more than one hundred species^31^.

Subsequent analyses identified two additional placozoan genera: *Hoilungia*^32^ and *Polyplacotoma*^33^. The nuclear genomes of both *T. adhaerens*^34^ and *H. hongkongensis* have been sequenced^32^, confirming distinct molecular features for each genus.

The overall morphology of *H. hongkongensis* was found to be very similar to that of *T. adhaerens*^32^, including the presence of the six major cell types^16^. A morphological study comparing different haplotypes of Placozoa (H1, H2, H5, H8 and H16 from different geographical origins^19^) also suggests that the basic bodyplan is conserved in placozoans, while subtle differences exist between clades.

In the present study, we used both transmission and scanning electron microscopy to characterize the cellular architecture in *H. hongkongensis*^32^ in more detail. While we did recognize the six basic cell types described both in *T. adhaerens*^16^ and *H. hongkongensis*^32^, we revealed additional populations of cells with distinctive morphological and positional features. These data corroborate the notion that the cellular diversity in placozoans is higher than anticipated.

## Material and Methods

### Culture

We worked with *H. hongkongensis* (mitochondrial 16S haplotype H13)^32^. Animals were cultured at 24±4°C in 9 cm-diameter Petri dishes containing artificial seawater (ASW; 35 ppm) with rice grains as food source^35^. For long-term maintenance, ASW was refreshed every 7 to 10 days (pH 8.0).

### Scanning electron microscopy (SEM)

To achieve fast fixation and preserve morphological features of placozoans in SEM, we used a protocol described elsewhere^36^. Briefly, one day before fixation, animals were transferred in a sterile Petri dish without food. Subsequently, we moved animals into 2 ml Eppendorf tubes with minimum ASW and added 2.5% glutaraldehyde (in 35 ppm ASW) for 60 min at room temperature. Next, the fixative was removed, and individuals were washed three times for 15 min in a 0.3 M NaCl and 0.05 M sodium cacodylate solution. Then this solution was gently removed, and we performed a secondary osmium fixation (0.3 M NaCl, 0.05 M cacodylate sodium, 1% OsO_4_) for 1 hour at 4°C. This solution was gently removed at room temperature. The specimens were dehydrated in acetone series (30%, 50%, 70%, 80%, 90%, 95%, 100%) with three 5-min washes. Animals were kept in 100% acetone at 4°C, and we performed critical point drying. The specimens were coated with platinum for 10 sec and examined under 15 kV using an FEI Quanta 250 scanning electron microscope (FEI, The Netherlands) at the Zoological Institute of Russian Academy of Science (Saint-Petersburg, Russia). In some cases, animals were broken with tungsten needles before platinum coating to reveal the interior structures. About 100 animals were used altogether.

### Transmission electron microscopy

Transmission electron microscopy (TEM) was performed exactly as described elsewhere on *H. hongkongensis* (H13)^32^ and *T. adhaerens* (H1)^16^. Briefly, animals were rapidly dipped in 20% BSA in ASW and placed in 0.1 mm-deep aluminum planchettes for high-pressure freezing (Wohlwend HPF Compact 02, Switzerland). Samples were processed by freeze-substitution (Leica AFS) for fixation (involving successive treatments in acetone with 0.1% tannic acid and 2% Osmium tetroxide) and Epon embedding. Polymerized samples were cut in 70-nm ultrathin sections collected on coated copper single-slot grids, which were then further contrasted and observed in a Philips CM100 transmission electron microscope.

### Light microscopy - differential interference contrast (DIC) microscopy

Light microscopic observations were made with a Nikon Ti2 fluorescent microscope equipped with a spinning disk and DIC optics.

## Results

### Gross morphology

Using scanning electron microscopy (SEM), we first examined the different surfaces of the placozoan *H. hongkongensis*. The lower epithelium (Fig. 1B) is the locomotory surface, and its high cell density results in a more densely ciliated surface as compared to the upper side of the animal (Fig. 1A), as described for *T. adhaerens*^11^. The shapes of cells on the upper epithelium are readily visible at high magnification, while on the lower surface, the high density of cilia largely masks the cells’ boundaries (Fig. 1C).

**Figure 1.**
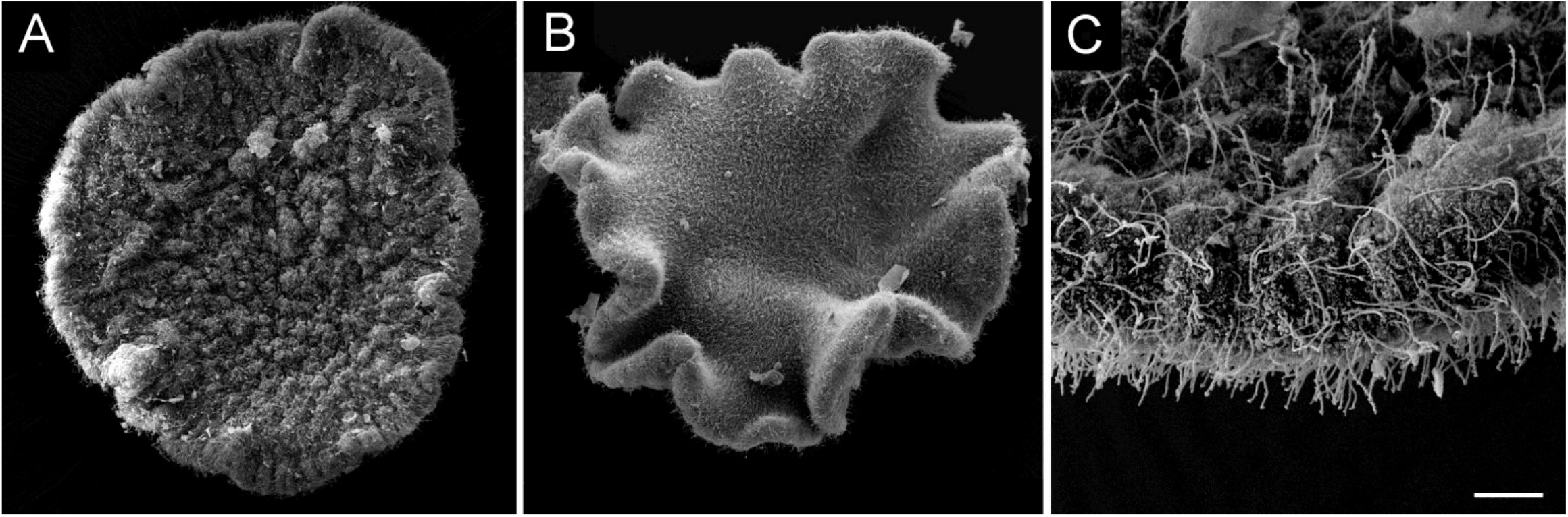
Morphology of *H. hongkongensis* (H13), viewed from the upper. (A) and lower (B) sides. Note that the animals were fixed while they were not adhering to a substrate, thereby showing great variability in body shapes, as they can form invaginations, ripples, curls, and can variably stretch and contract. (C) Ciliated epithelia in *H. hongkongensis* (H13). The lower surface is more densely ciliated than the upper surface. Some cilia appear very long and may belong to distinct cell types. Scale bar: 50 μm in A, B; 8 μm in C.

The gross morphology of the placozoan *H. hongkongensis* had been shown to be similar to that of *T. adhaerens* using transmission electron microscopy (TEM)^32^. When we inspected fractioned freeze animals by SEM, we were able to view the internal organization of the animal (Fig. 2A). While the lower epithelial layer is thick - as it is composed mainly of densely packed columnar cells of different heights - the middle layer is overall loose and comprises very large cells with extensions. Most cells in the upper epithelium form a flat, thin surface, but other cell types can be observed as well, which will be described below. We then extended the analysis by imaging at the TEM entire cross-sections of *H. hongkongensis* (Fig. 2B). These images corroborate a three-layer bodyplan, with two epithelial layers that encompass a more loose middle layer. Also easily recognized are large osmiophilic vacuole-like structures corresponding to Schulze’s shiny spheres (3–5 μm), on the entire upper side of the animal (Fig. 2B, B1). They belong to a population of cells that has not been characterized yet (see below). In turn, somewhat smaller osmiophilic vacuole-like inclusions (2-3 μm), corresponding to Schulze’s matte-finished spheres, are distributed in the medial part of the lower epithelium, and are close to the lower surface of the animal (Fig. 2B, B2). They have readily been identified as the most distal vacuole of lipophil cells^16^. At the TEM, both shiny and matte-finished spheres are lipoid in nature and show varying degrees of affinity to osmium (Figs. 2, 3, 6), possibly depending on the preparation procedures or physiological state of the animal. Whether their content - which is not known so far - is secreted by the animal and whether the two cell types producing these structures are related is unclear. Of note, one can observe at higher magnification that adherens junctions link all cells of the epithelia. The vast majority of these cells possess a single cilium and microvilli-like organization (Fig. 2C;^32^, which was similar to a system of folds from the columnar cells in *T. adhaerens*^8,37^; and best described as a “spongy meshwork of fenestrated folds” apparently distinct from microvilli (see details in^13,16,38^). In *H. hongkongensis*, a pronounced glycocalyx-like structure can be seen on the surface of the lower epithelium (Fig. 2C).

**Figure 2.**
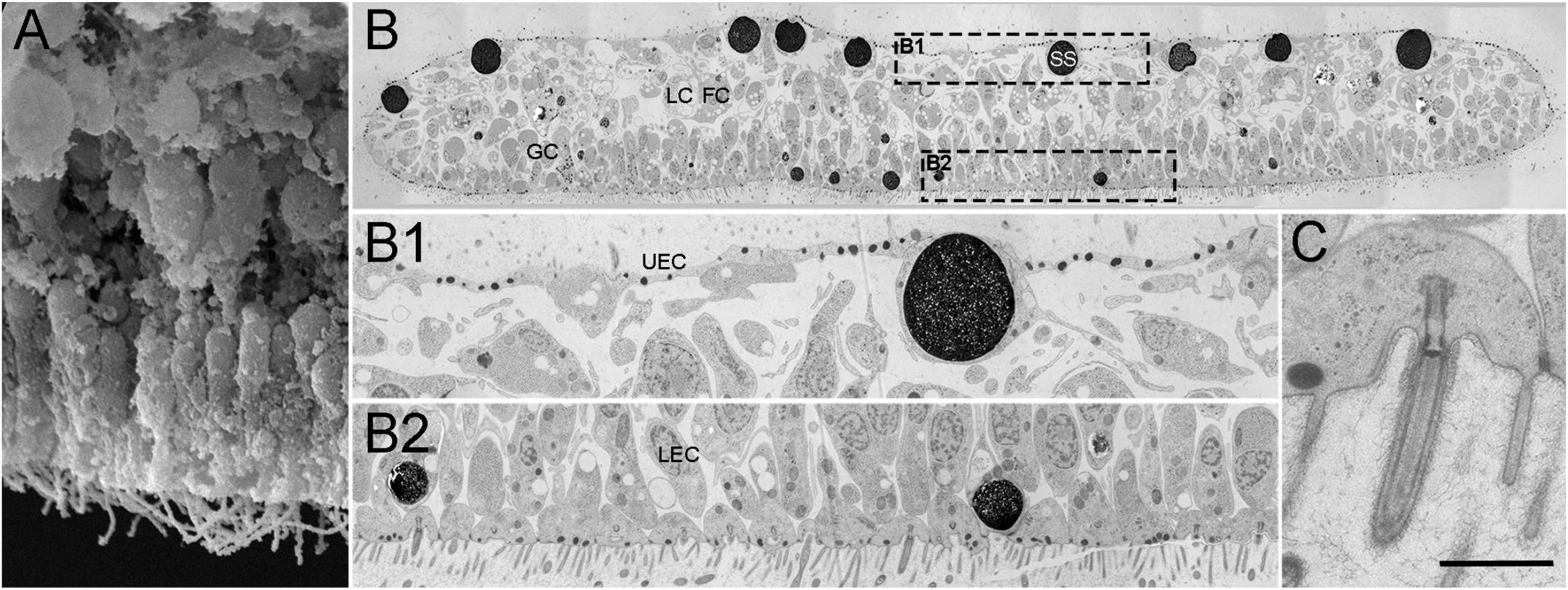
Gross anatomy of *H. hongkongensis* (H13). (A) A SEM micrograph from the animal’s inside reveals, from top to bottom, the flat cells of the upper epithelium, the bulky cells of the middle layer, and the numerous, mostly elongated cells of the lower epithelium. Note that micro-cavities were often observed between groups of densely packed cells of the middle and lower layers. (B) At the TEM, a full cross-section micrograph of a representative animal (approximately 250 μm-wide and 30 μm-thick) shows the three-layer bodyplan and upper and lower epithelia merging at the edges. Framed regions of the upper and lower epithelia are shown at higher magnification in (B1, B2) and readily reflect cell diversity. In B1, the shiny sphere-containing cell shows an elongated process toward the middle layer and protrudes between two canonical flat epithelial cells. In B2, the vast majority of columnar cells are ciliated. In contrast, lipophil cells are not ciliated, and their outermost granule (“matte-finished sphere”) about the surface. As illustrated in (C) at the level of the lower epithelium, all cells are joined by adherens junctions, and most bear a cilium and microvilli-like structure (see text for details). Note the ciliary root and the ciliary pit, which can be more or less deep in different cell types, as well as some unidentified glycocalyx-like material outside the membrane. FC, fiber cell; GC, gland cell; LC, lipophil cell; LEC, lower epithelial cell; UEC, upper epithelial cell; SS, shiny sphere. Scale bar: 6 μm in A; 18 μm in B; 5 μm in B1, B2; 1 μm in C.

### Lower epithelium

In *H. hongkongensis* as in *T. adhaerens*, the lower epithelium is mostly composed of columnar lower epithelial cells and lipophil cells (Figs. 2B2, 3E;^32^ Both cells play roles in external digestion and uptake of nutrients at the basal surface. This process is probably coordinated by different types of gland cells embedded in the lower epithelium^13,16,20–22^).

**Figure 3.**
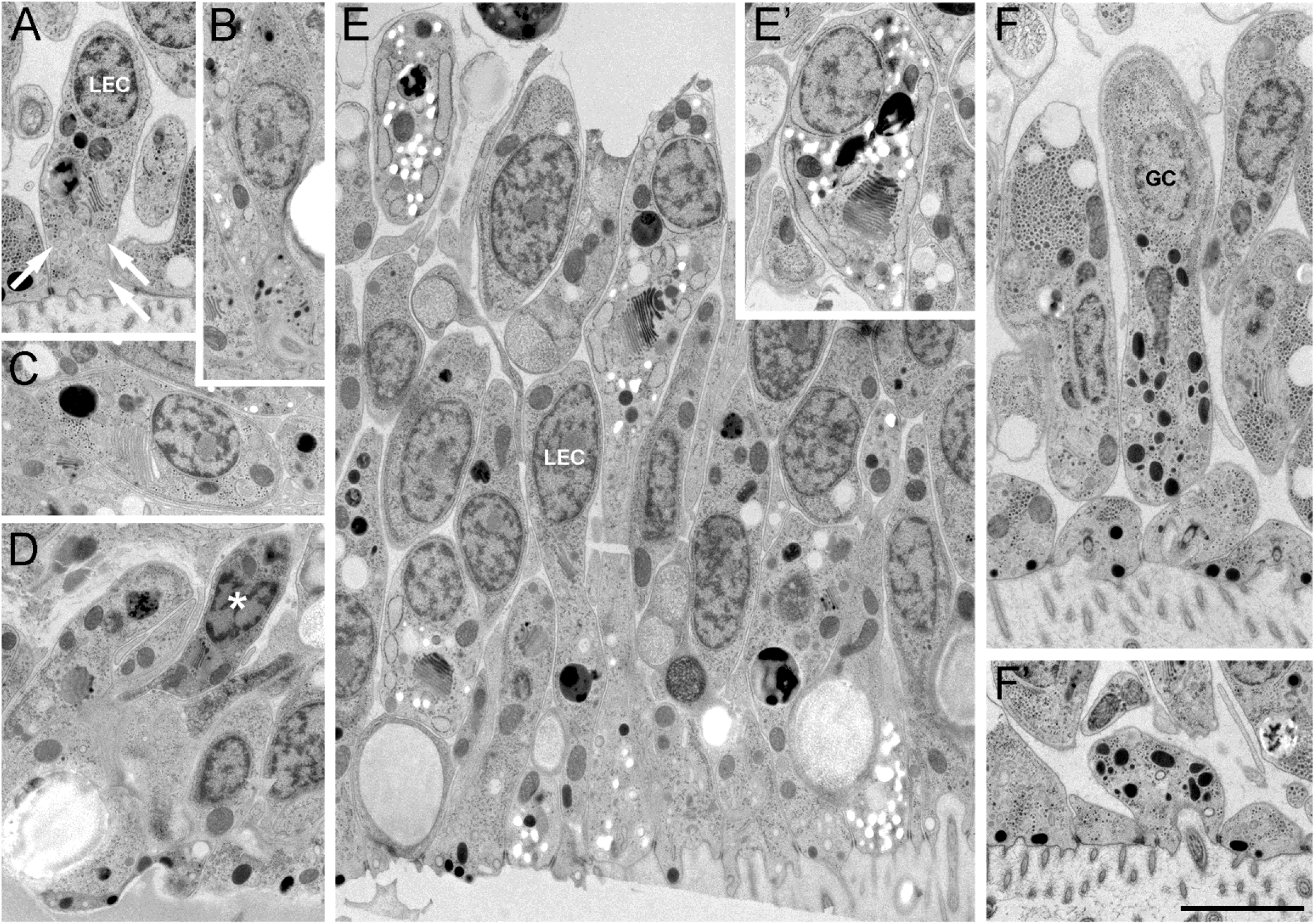
Cell diversity in the lower epithelium of *H. hongkongensis* (H13). (A) Classical lower epithelial cells are columnar and polarized. They often bear clusters of grey-ish vesicles, possibly reflecting pinocytic events (arrows). (B) Small polarized, ciliated cells present clusters of small dark vesicles near the lower surface of the animal; they likely correspond to an undescribed population of secretory cells. (C, D) Cells with specific content (C) or with a dark appearance (asterisk) are regularly spotted among the cells of the lower epithelium; they may not reach the surface. (E, E’) In some areas of the epithelium, among lower epithelial cells and lipophil cells, a population of polarized cells contain clusters of very clear vesicles close to the Golgi apparatus or reaching the surface; they may correspond to mucus-secreting cells. (F, F’) Typical gland cells – ciliated, polarized secretory-like cells with large dense vesicles – are present at the inner rim of the lower epithelium. GC, gland cell; LEC, lower epithelial cell; Scale bar: 3 μm.

Lower epithelial cells are highly polarized in *H. hongkongensis*, and often contain numerous pinocytic-like vesicles at their basal pole (Fig. 3A, E). Lipophil cells – the second most abundant cell type – have somata larger and taller than those of epithelial cells and reach deep into the middle cell layer (Figs. 2A, 3E, 5A). At the SEM, we noted that lipophil cells, together with other cells, delineated numerous microcavities (areas without cells or their processes), which were located in the middle layer between different cell types (Fig. 2A). These microcavities can be seen in SEM micrographs of *T. adhaerens* as well^16^; whether they have a role remains elusive, however.

Several other cell types were observed scattered at low density across the lower epithelium in *H. hongkongensis* (Fig. 3B-F) and reaching the basal surface. Ciliated cells with middle-sized, irregularly shaped, dark granules (Fig. 3F) likely correspond to the gland cells described in *T. adhaerens*^16^. In *H. hongkongensis*, these vesicles are smaller and more irregular, so they might correspond to different cell types.

Apparently, one type of gland cells is found mostly in the rim area of the animal, while another type is found throughout the lower epithelium^22^. Other ciliated cells contained tiny dark granules (Fig. 3B). We also came across long, highly polarized cells with a large Golgi and numerous clear granules accumulating at their basal pole, microvilli, and no cilium (Fig. 3E, E’). These cells might be similar to mucin-releasing cells recently described in *T. adhaerens*^22^ and/or the gland cells without cilium described by Grell and others (summarized in Grell and Ruthmann, 1991). Several additional cells, which can probably be regarded as other cell types, were found embedded in the lower epithelium (and no apparent contact to the outside), such as cells whose cytoplasm is enriched in granular material (Fig. 3C) and cells with a very dark cytoplasm (Fig. 3D).

Of note, we observed on the water-facing side of the lower epithelium, regularly spaced small pores/recesses (approx. 1 μm in diameter; not shown). These might be sites of the release of different secretory components from lipophil (e.g., be the point where the large vacuoles of lipophile cells lie), mucin-releasing, or other secretory cells.

Lipophil cells have gotten their name from their numerous, large, lipid-rich vacuolar structures^16^. As illustrated in Fig. 4, lipophile cells are sturdy (A, B), elongated (C) or grossly ramified. In fact, we found that lipophil cells are highly variable in size and shape, and it is conceivable that different stages or even subtypes of lipophil cells exist. Also, their shape may depend on the local environment/cellular configuration as lipophil cells are often observed near fiber cells in the middle layer (Fig. 4B, 5A).

**Figure 4.**
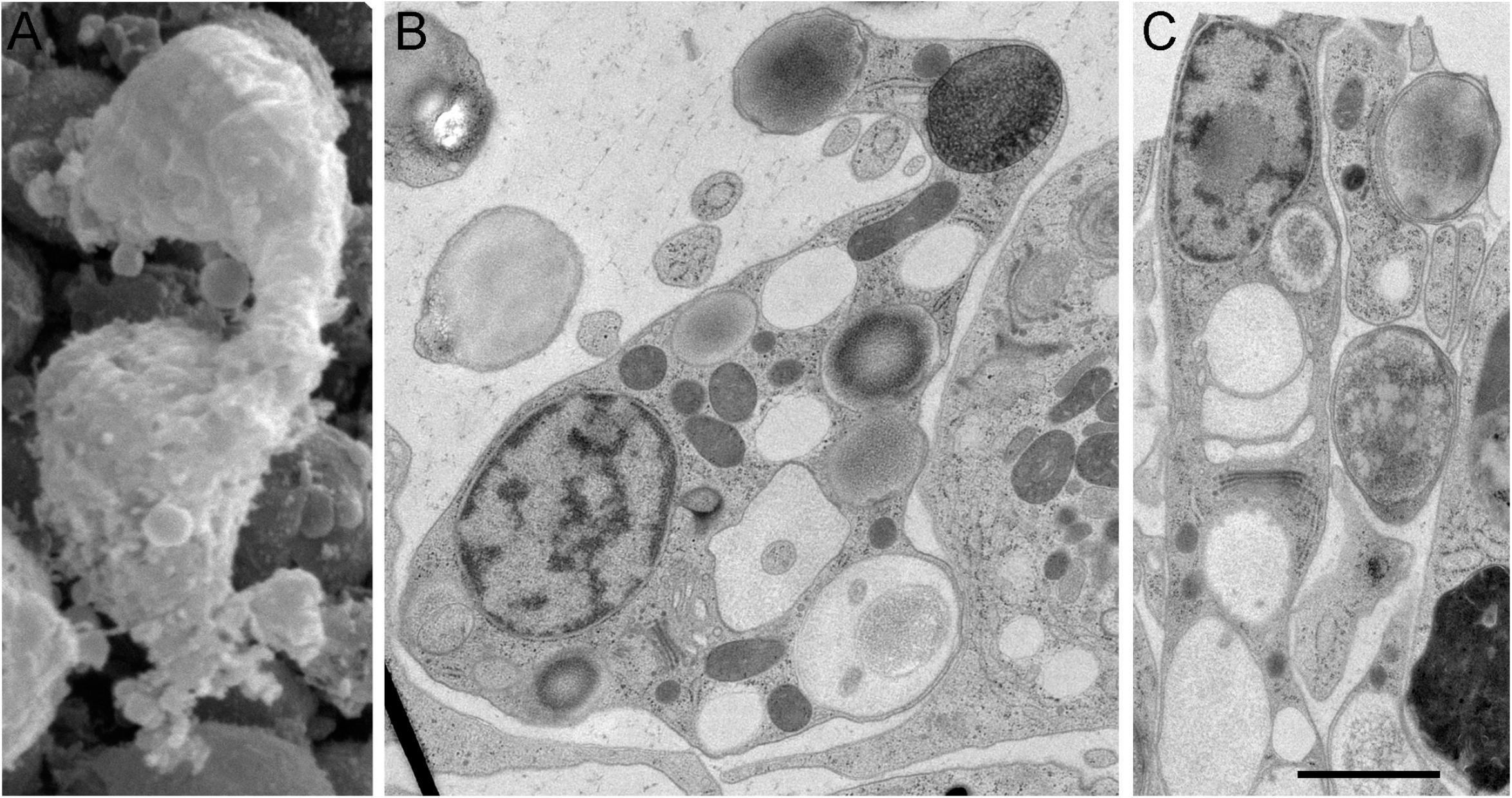
Ultrastructure of lipophil cells in *H. hongkongensis* (H13). (A) At the SEM, lipophil cells are at times seen bulging from the lower epithelium in the middle layer, often in close vicinity to fiber cells. Their shapes are usually elongated (in the lower epithelium, where they are interspersed between lower epithelial cells (C) or more complex depending on their cellular environment (B) and position in the animal. In all cases, they exhibit large, heterogeneous vacuoles - rather clear for these close to the trans-Golgi network, otherwise with various shades of grey (i.e., affinity to osmium) or complexity, as some of the vacuoles contain membranous material. Their most basal vacuole (matte-finished sphere), abutting the lower surface, is often electron-dense. Scale bar: 2 μm in A; 1.5 μm in B, C.

**Figure 5.**
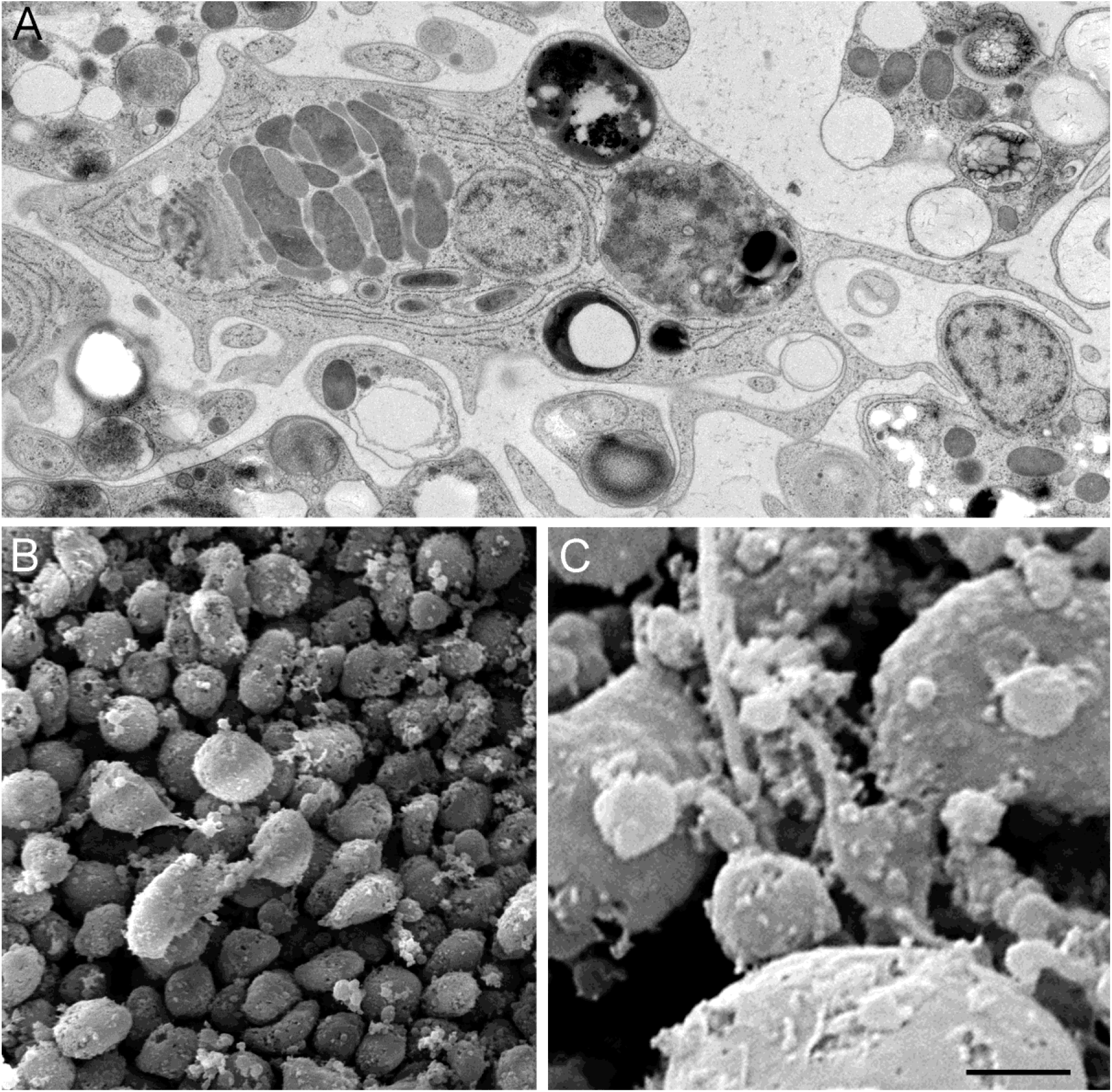
Ultrastructure of cells in the middle layer of *H. hongkongensis* (H13). (A) Each fiber cell has a bulky soma and numerous thin processes; complex, heterogenous vacuoles in varying numbers; a single large mitochondrial cluster; and bacteria inside the endoplasmic reticulum. Structures reminiscent of exosomes or vacuoles from lipophil cells are present around fiber cells. (B) On this SEM micrograph, a group of four fiber cells lie on top of cells of the lower epithelium, from which lipophil cells are emerging; fiber cells are interconnected, and their processes reach deep between epithelial cells. (C) Unknown small cells with processes were observed in the close vicinity of fiber cells. Scale bar: 1.5 μm in A; 5 μm in B; 1.5 μm in C.

### Middle layer

In *T. adhaerens*, the middle layer of the animal is quite loosely packed and consists mostly of fiber cells. Fiber cells are very large cells with uniquely branched cellular extensions and processes^9^. Their ramification pattern suggests that they contribute to the connectivity of all layers, forming a potential network supporting and capable of coordinated reactions and, perhaps, behavioral integration in general^13,16^.

In *H. hongkongensis*, fiber cells were also easily identified in the middle layer, with their distinct shape and sub-cellular features (Fig. 5A, B). They are large (the soma is 5-15 μm wide), elongated, and asymmetric cells with numerous processes mingling with lipophil cells and reaching laterally and deep in the upper and lower epithelia (Figs. 2B, 5A). They possess a large nucleus, a characteristic mitochondrial complex (a bulky cluster of alternating layers of mitochondria and vacuoles with unknown function), large vacuoles with complex, heterogeneous content (named concrement vacuoles in *T. adhaerens*^13^), and a well-developed endoplasmic reticulum (Fig. 5A) in which bacteria are present. Genomic analysis was made in *Trichoplax* sp. (H2)^39^.

Occasionally, smaller cells were observed in the vicinity of fiber cells. They had a star-like shape with 1 to 3 thin, long processes (less than 1 μm in diameter and 10 to 50 μm in length; Fig. 5C). Such cells have so far not been reported in *T. adhaerens*.

### Upper layer

By SEM, the outer upper surface of the animal appeared as a mosaic of large polygonal cells bearing a single cilium, scattered with dome-shaped elements (Fig. 6A). The large contours correspond to upper epithelial cells (Fig. 6B): ciliated, T-shaped cells known to form a thin epithelium, described for *T. adhaerens*^13,16^, and observed here in *H. hongkongensis* (Fig. 6B, C). They were connected with each other by adherens junctions, and contained numerous dark granules close to the surface that likely correspond to pigment granules, which were observed by DIC on living preparations (not shown) and were reported in *T. adhaerens*^13,16^.

**Figure 6.**
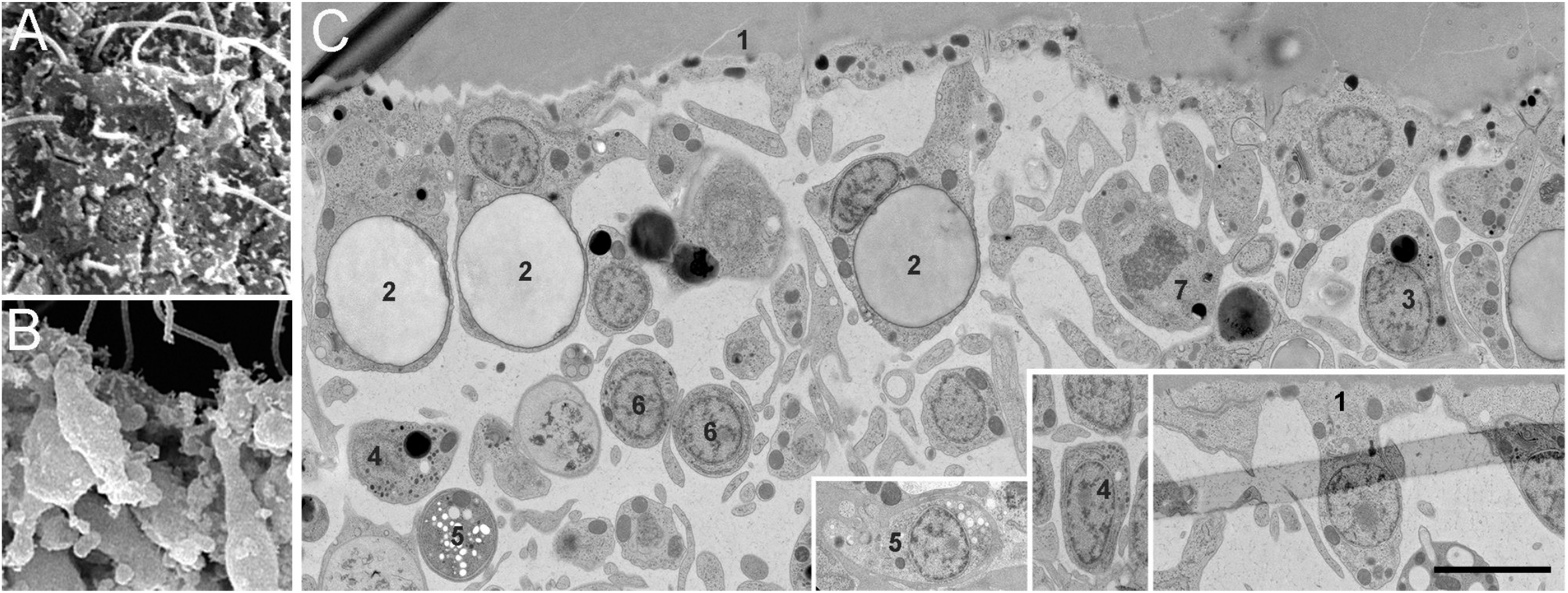
Cell diversity in the upper epithelium of *H. hongkongensis* (H13). (A) At the SEM, contours of cells composing the upper epithelium can easily be made out. Most of the cells are ciliated. Some concave discs can be seen, which may correspond to shiny spheres abutting the surface. (B, C) Cells from the upper epithelium are essentially bound to other cells at their basal pole (toward the upper surface) by adherens junctions, and the main cell body (including the nucleus) is sunk toward the inside of the animal. They may receive local and maybe transient contact from other cells, such as fiber cells. Canonical upper epithelial cells (“1”) are elongated T-shaped ciliated epithelial cells with pigment granules. They have a large flattened surface area, are not ramified, and their nucleus occupies most of the cytoplasm. Towards the upper surface, they contain electron-dense inclusions. (C) A variety of other cell types can be identified in the upper epithelium and directly underneath. Some cells (“2”), ‘sphere cells’, have a large inclusion that might correspond to a shiny sphere; they have no pigment granules, a small nucleus and a few processes; some cells (“3”) contain small vesicles, reach the surface via a narrow neck and have no cilium; right underneath the cells forming the surface, populations of small putative secretory cells with numerous small dense vesicles (“4”) or small clear vesicles (“5”) are observed; small rounded cells are seen in pairs (“6”) that may have recently duplicated; other cells, reminiscent of macrophages with an atypical nucleus and numerous ramifications are also observed (“7”). Processes of fiber cells are omnipresent. Scale bar: 5 μm in A; 4 μm in B; 3 μm in C.

In *T. adhaerens*, shiny spheres in the upper epithelium have been described early on^8^, but were not observed in all studies^16^. According to earlier observations^19^ and based on our new results on *H. hongkongensis* the nature and origin of shiny spheres was misdescribed as they were often considered as “independent” organelles. In our observed specimens, shiny spheres are frequently encountered and corresponded to large vacuoles belonging to a specific population of cells (see also^19^). These sphere-bearing cells differed from regular upper epithelial cells as they did not have a large outer surface and sent a few ramifications toward the inner cavity. The dome-shaped structures seen by SEM at the animal’s surface (Fig. 6A) may correspond to the shiny spheres abutting the surface (Fig. 2). This confirms a previous assumption by Guidi et al. (2011), who described the remnants of released shiny spheres as ‘concave discs’. Our results add a new aspect to the conclusions drawn by the aforementioned study. The position of a shiny sphere within ‘sphere cells’ (close to the outer surface or deeper inside) as well as the sphere’s density may correspond to different maturation states of the cell or even different physiological states of the animal.

Within the upper epithelium (Fig. 6C), a few cells bearing clusters of small dense granules reached the surface via a narrow neck. They may represent a specific population of secretory cells. Directly underneath the epithelium, several cell populations with distinct morphological features were observed, which to our knowledge, have never been previously described. In contrast to the lower epithelium, cells were loosely packed, and a clear border to the fiber cell layer could not be recognized. Yet many elongated processes were observed, suggesting that cells in this area, while not tightly opposed, may connect to each other and/or with fiber cells. No specialized subcellular features were observed at these contact points that could reflect a specialized form of local (e.g., chemical or electrical) communication.

These newly identified cell types varied in size, shape and cellular content. Putative secretory cells were singled-out, which contain clusters of clear vs. electron-dense vesicles of different sizes, suggesting different contents. In each cell type, granules’ population was homogenous. Often, cells with similar subcellular characteristics were close to each other, suggesting that the cells divided and likely differentiated on the spot. Atypical cells, remotely resembling immune system-related cells, were also observed (Fig. 6).

On the inside, near the upper edge, crystal cells were identified in *T. adhaerens*^16^. These spherical cells contain a central aragonite crystal and have a flattened, laterally located nucleus^18^. They are thought to function as gravity sensors^18^. Crystal cells with similar morphology were observed in *H. hongkongensis*^32,40^. As in other placozoans, crystal cells were located under the upper epithelium and did not have any noticeable direct contact with the external environment (Fig. 7A). Due to their birefringent crystal, they can be readily observed using DIC microscopy (Fig. 7B). In *H. hongkongensis*, crystal cells were quite scarce and usually spread within a concentric band along the marginal zone.

**Figure 7.**
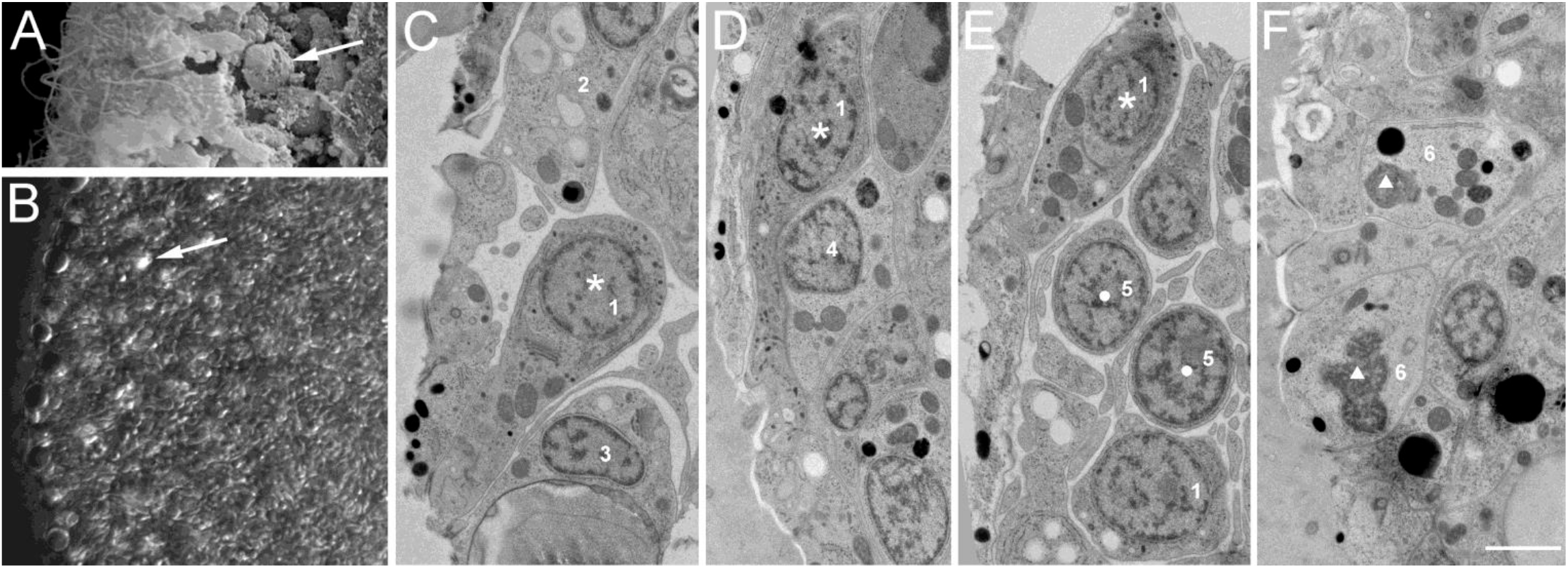
Cell diversity in the marginal zone of *H. hongkongensis* (H13). (A) Crystal cells (arrow) were rather scarce and located at the marginal edge of animals under the upper epithelium, sometimes close to the rim. (B) Observation of a live specimen under DIC illumination illustrates the numerous cilia visible at the edge, large vacuoles of cells present at the periphery, and a bit toward the inside, the typical birefringent crystals of crystal cells (arrow). (C-E) At the TEM, cells with a dark cytoplasm, numerous irregularly-shaped dark granules, a long neck reaching the surface, and a cilium were frequently observed (asterisk). (E) some round cells with little cytoplasm might correspond to proliferative cells (dot). (F) Atypical cells with a poorly defined nucleus, numerous mitochondria, and large dense-core vesicles are also observed (triangle). At least 6 different morphologically distinct cell types (numbered 1-6) can be distinguished in this figure. Scale bar: 6 μm in A; 16 μm in B; 1.5 μm in C-F.

### Marginal zone

As mentioned earlier, the lower epithelium accommodates at its rim concentric rings of gland cells and mucocytes^22^. This zone was also found to contain different populations of peptidergic cells^20,21^, suggesting that different gland cells that secrete different signaling molecules exist.

In the marginal zone, which one can define as the outermost rim of the animal and the area where upper and lower epithelia meet (Fig. 2B), small ovoid-like cells were described in *T. adhaerens*^19^ This scantily described area has been presented as a putative proliferative zone^41^, and is thought to contain freshly-dividing cells^41,42^.

In *H. hongkongensis*, we observed in the marginal zone elongated cells with a narrow neck and opening to the surface and a cilium and no microvilli, which contained very small dark granules (Fig. 7C-E). They likely belong to yet another population of sensory or secretory cells. Small-size cells with a round nucleus and little cytoplasm were frequently observed inwards, near these cells (Fig. 7D). They did not seem to reach the surface. They might correspond to the population described by Guidi et al.^19^ and reflect a proliferative zone. Last, atypical cells, similar to these observed under the upper epithelial surface, were present in this region (Fig. 7F).

## Discussion

The placozoan *T. adhaerens* has a very simple shape, this of a small plate, with an upper (protective) and a lower (nutritive) epithelial layer encompassing a loose middle layer. While not feeding, it presents a constantly changing outline due to its continuous movements and body contractions. As the simple bauplan of *T. adhaerens* did not correspond to any of the other basic lineages of animals – sponges, cnidarians, ctenophores (commonly known as comb jellies), and bilaterians – the animal was placed into its own phylum Placozoa^15^. The exact phylogenetic position of the phylum Placozoa is still debated, although a consensus seems to emerge that views Placozoa as the sister group to the clade Cnidaria and Bilateria^32,43–46^. In recent years, different placozoans species have been discovered, starting to reveal the genetic diversity of the phylum^26,27,31,33,47^.

One generation ago, these simple animals were thought to have only four types of cells^13^. Yet over the last years, several morphological studies have unraveled several additional major cell types^16,48^. A recent single-cell sequencing study has indicated the existence of an even larger variety of cell states, which might support the existence of additional cell (sub)types^3^. Indeed, it turned out that the lower epithelium, which is specialized for extracellular digestion, contains different types of gland cells and mucin-secreting cells^16,22,48^. Moreover, the complex and rapid movements, action potentials, and coordinated food uptake of the animal are controlled by integrative systems, including several different types of endocrine cells, suggesting that the animal is much more complex than previously assumed^21,49–53^.

Here we have taken a closer look at the morphology and cell types of *H. hongkongensis* (H13 haplotype from the South China Sea), which, based on comparative genomics, belongs to a different genus of placozoans but has a very similar basic bauplan as *T. adhaerens*^32,47^. We wanted to describe its morphology in more detail and also search for additional cell types in this species.

Ours and other researcher’s longterm goal is to match the cell types found by single-cell sequencing approaches^2,3^ to cell types established on morphological grounds. This will serve as a basis to understand the function of the different cells and how the cells sense their surroundings and communicate with each other to serve food uptake, to trigger movements, and to control other behaviors of placozoans. Addressing this will help shed light on the intriguing question of whether the different cell types and, by extension, the three cell layers in placozoans are homologous or not to cell types and germ layers found in the other major animal lineages^2,3,54^.

It appears that several specialized cell types, regarded as the “basal building blocks of multicellular organisms”, have already emerged in placozoans. For example, it is being discussed whether the cell types in the nutritive epithelium of placozoans resemble these of the intestine^4,55^. Likewise, it needs to be explored whether (neuro)peptide-releasing cells discovered in placozoans are homologous to neuroendocrine or even neuron-like cells^21^.

Here, we corroborated the notion that the overall bodyplan and cell types of *H. hongkongensis* are very similar to that of *T. adhaerens*^32^, suggesting that most findings apply to both species. Nevertheless, subtle morphological differences probably exist and need to be further investigated. One morphological difference is the somewhat different position of crystal cells in the two species. Crystal cells are sparse and located underneath the upper epithelium in both genera. They are thought to function as gravity sensors^18^.

Several other cell types have been already described morphologically, and functions have been ascribed to them, as we will outline below. Moreover, we found ultrastructural hints for additional cell types for which currently no functional description is available.

The lower nutritive epithelium is formed by ciliated columnar epithelial cells that drive the animals’ gliding through ciliary beating. These cells may also be involved in the uptake of nutrients^8,37,56^ after release of digestive enzymes for external digestion by the large lipophil cells^16,22,48^, which are interspersed in the epithelium. However, to what extent the digestive system of placozoans is homologous to that of other animals is not clear yet. The cell bodies of lipophil cells reach deep into the middle layer. Eitel et al. (2018) and our study in *H. hongkongensis* show that the basic morphology of lipophil cells is comparable to those described in *T. adhaerens*. Yet we report a remarkable heterogeneity of these cells, which vary in size, position, and shapes. A subset of lipophil cells might not reach the outer epithelial surface and, therefore, differ in their secretion sites. The functional and structural diversity of lipophil cells has not been investigated yet in *T. adhaerens*, and it would be interesting to identify the molecular markers for different putative subpopulations or different functional states of these cells.

In *T. adhaerens* as in *H. hongkongensis*, secretory-like cells corresponding to canonical gland cells are scattered throughout the lower epithelium^16,32^, which are possibly involved in local regulatory processes. Recently, two different populations of ciliated gland cells were found in *T. adhaerens*^22^. One population was found near the edge of the animal, whereas the other was found more centrally. In addition, mucocytes without a cilium were identified^22^, concentrated close to the edge of the animal but are also found in the central region of the lower epithelium. Mucus secretion supports adhesion of the animals^22^ In our study, we have established that similar cell types exist in *H. hongkongensis* as well and have extended their morphological description. Note that these cells morphologically resemble cells of the digestive and respiratory tracts of both vertebrates and invertebrates^57–62^.

Both in *T. adhaerens* and *H. hongkongensis*, the middle layer consists mainly of fiber cells^8,12,63^. They are large, have long extensions, and contain a cluster of mitochondria and complex vacuoles. Fiber cells, possibly, form a syncytium^9^ that connect and mediate an integration across all other cell types. Hence, fiber cells may support systemic organismal functions and complex behaviors. Occasionally in *H. hongkongensis*, we came across smaller star-shaped cells with thin, long processes. However, more research is needed to determine whether they are merely smaller fiber cells or constitute a different population of cells.

Fiber cells of all placozoan lineages studied so far (including *H. hongkongensis*) possess a rickettsial endosymbiont inside of the endoplasmic reticulum^8,19,39,64,65^. Based on bacterial 16S sequencing, the bacteria observed in fiber cells of *H. hongkongensis* belong to a different *Aquarickettsia* species than the *T. adhaerens* endosymbiont^66^. No signs for host-specificity were identified that would imply close co-evolution of a specific bacterial species and its host. Of note, in *T. adhaerens sp*. (16S haplotype H2), an unrelated bacterium (Candidatus *Ruthmannia eludens*) was also identified in lipophil cells^65^. Neither our study on *H. hongkongensis* nor previous analyses on *T. adhaerens* (H1) have identified bacteria in lipophil cells. The putative absence in other placozoans, as well as the presence of versatile biosynthesis pathways^65^ in *Ruthmannia* might, therefore, suggest a parasitic rather than a symbiotic lifestyle of this bacterial species. Future molecular studies on a range of placozoan species will help to elucidate the diversity, distribution and function of the two placozoan-associated bacterial lineages known to date.

The upper epithelium of *T. adhaerens* has been described as a homogenous population of flat, ciliated cells forming a protective layer. Their cell bodies with a nucleus hang from the flat upper part, a structure that has been described as T-shape^13,16^. Very similar epithelial cells form the upper epithelium in *H. hongkongensis*, but we found several other cell types to be present in and underneath the upper epithelium. One type of cell contains shiny spheres, as described above. Note that shiny spheres were not observed in the laboratory *T. adhaerens* strain used by Smith and colleagues^16^, but have been described by Grell and others^13^. Presence or absence of shiny spheres within the same strain in conjunction with the presence/absence of concave discs in various placozoan strains^19^ indicates that hidden developmental and/or seasonal stages might exist in placozoans.

The role of shiny spheres remains enigmatic still. It has been suggested that these structures are part of a chemical defense mechanism, which may use paralytic substances toxic for predators of placozoans^67^. As this lipid-rich organelle is not a non-cellular lipid inclusion as suggested earlier^13^ but is produced by a distinct cell, it should be possible in the future to identify this cell type from its expression profile. Another intriguing matter is whether the cells that produce shiny spheres in the upper epithelium, we here term ‘sphere cells’, and lipophil cells, which generate morphologically similar but smaller matte-finished spheres at the lower surface, are related. Do both cell types extrude the content of their large vacuole-like structures into the extracellular medium, or do these prominent structures serve different functions? Some of the cells with larger vacuole-like structures were underneath the upper epithelium but did not reach the upper surface. Whether these cells might correspond to an immature form of shinysphere-containing cells, which would further mature and move into the upper epithelium remains to be clarified.

In our study, we came across putative secretory cells in the lower and the upper epithelium and other cells that are located in the marginal zone. These secretory cells could correspond to one of the several cell populations that contain different secretory signaling peptides ([neuro]peptide-like molecules). About ten different (neuro)secretory cells are located in distinct regions of both epithelial and in the inner layers of *T. adhaerens*^21^. Interestingly, many of these lower-frequency cell types are organized in separate concentric circles in the different layers of the animal. They may be rather difficult to tell apart based solely on their morphology, and it is possible that different secretory cells are currently lumped together. For example, different secretory-like cells containing the predicted prohormones for endomorphin (also referred to as YPFF)^20^, YYamide, and FFNPamide, and cells containing the insulin-like prohormone^68^ have been found in the lower epithelium, while other peptidergic cells are in the upper epithelium (SITFamide) and marginal region (SIFGamide)^21^. In order to distinguish them on an ultrastructural level, specific probes are required. Nevertheless, together with recent microchemical data about the low molecular weight transmitter candidates^69,70^, the repertoire of secretory cells shows that placozoans are able to release a variety of signaling molecules that probably elicit different behaviors of the animal. The target cells for the different peptides and other intercellular messengers, including nitric oxide^71^, need to be identified yet.

Besides clarifying which basic cell types and derivatives exist, where they are located in the body, and if they do belong to the basic makeup of all placozoans, a formidable challenge will be to elucidate what the precise function of each cell type is and how the different cells communicate and coordinate the behavior of the animal. Does most intercellular communication occur mostly by paracrine signaling or do many cells connect to other cells via longer extensions as it has been shown for fiber cells? The loose packing and the many cellular protrusions in the interior of the animal will require three-dimensional reconstructions to shed light on the potential connectivity of the cells and the organization of the bodyplan.

## Data Availability

The data that support the findings of this study are available on https://zenodo.org.

## Author contributions

D.Y.R., F.V., D.F. and L.L.M. designed the study; D.Y.R., F.V., J.D., M.A.N., M.E. and L.L.M analyzed the data; D.Y.R., L.L.M, and D.F. wrote the paper; and all authors reviewed, commented on, and edited the manuscript.

## Acknowledgments

We thank E. Bedoshvili, A. Miroliubov, and V. Starunov for their help in electron microscopy, sample preparation, and advice for EM protocols. This work was supported by the Human Frontiers Science Program (RGP0060/2017) and National Science Foundation (1146575, 1557923, 1548121, and 1645219) grants to L.L.M.; Russian Ministry of Science and High Education (agreement 075-15-2020-801) grant to D.R. and the Swiss National Science Foundation (#31003A_182732) grant to D.F. The research reported in this publication was also supported in part by the National Institute of Neurological Disorders and Stroke of the National Institutes of Health under Award Number R01NS114491 (to L.L.M.). M. Eitel acknowledges funding from the European Union’s Horizon 2020 research and innovation program under the Marie Skłodowska-Curie grant agreement No 764840. The content is solely the responsibility of the authors and does not necessarily represent the official views of the National Institutes of Health.

